# Decoding of human hand actions to handle missing limbs in Neuroprosthetics

**DOI:** 10.1101/015040

**Authors:** Jovana J. Belić, Aldo A. Faisal

**Affiliations:** Department of Bioengineering, Imperial College London, London, UK; Faculty of Electrical Engineering, University of Belgrade, Belgrade, Serbia; Department of Computational Biology, Royal Institute of Technology KTH, Stockholm, Sweden; Department of Computing, Imperial College London, London, UK; MRC Clinical Sciences Center, London, UK

**Keywords:** Neurotechnology, motor control, prosthetic hand, neuroprosthetics, movement variability, Bayesian classifier, PPCA, activities of daily living, finger movement

## Abstract

The only way we can interact with the world is through movements, and our primary interactions are via the hands, thus any loss of hand function has immediate impact on our quality of life. However, to date it has not been systematically assessed how coordination in the hand’s joints affects every day actions. This is important for two fundamental reasons. Firstly, to understand the representations and computations underlying motor control “in-the-wild” situations, and secondly to develop smarter controllers for prosthetic hands that have the same functionality as natural limbs. In this work we exploit the correlation structure of our hand and finger movements in daily-life. The novelty of our idea is that instead of averaging variability out, we take the view that the structure of variability may contain valuable information about the task being performed. We asked seven subjects to interact in 17 daily-life situations, and quantified behaviour in a principled manner using CyberGlove body sensor networks that, after accurate calibration, track all major joints of the hand. Our key findings are: 1. We confirmed that hand control in daily-life tasks is very low-dimensional, with four to five dimensions being sufficient to explain 80-90% of the variability in the natural movement data. 2. We established a universally applicable measure of manipulative complexity that allowed us to measure and compare limb movements across tasks. We used Bayesian latent variable models to model the low-dimensional structure of finger joint angles in natural actions. 3. This allowed us to build a naïve classifier that within the first 1000ms of action initiation (from a flat hand start configuration) predicted which of the 17 actions was going to be executed - enabling us to reliably predict the action intention from very short-time-scale initial data, further revealing the foreseeable nature of hand movements for control of neuroprosthetics and tele operation purposes. 4. Using the Expectation-Maximization algorithm on our latent variable model permitted us to reconstruct with high accuracy (<5**°**-6**°** MAE) the movement trajectory of missing fingers by simply tracking the remaining fingers. Overall, our results suggest the hypothesis that specific hand actions are orchestrated by the brain in such a way that in the natural tasks of daily-life there is sufficient redundancy and predictability to be directly exploitable for neuroprosthetics.

## 1. Introduction

The human hand is a highly complex actuator and perhaps the most important and diverse tool we use to interact with the environment. The hand is capable of both a powerful grip to push, pull, or twist objects, and a precise grip to twist and turn small objects or handles (Napier, 1980). These are just a few of the countless gestures we can use and learn. Anatomically, the hand comprises a total of 27 bones, 18 joints, and 39 muscles (Tubiana, 1981), which afford over 20 degrees of freedom (DOF) (Stockwell, 1981; Soechting and Flanders, 1997; Jones and Lederman, 2006). The number of degrees of freedom is an important characterization of the human hand because it defines the dimensionality of the control problem that has to be solved by the motor system. However, previous studies of human motor control showed that normal hand behaviour uses only a small subset of possible hand configurations (Todorov and Ghahramani, 2004; Weiss and Flanders, 2004; Ingram et al., 2008; Valero-Cuevas et al., 2009). It is known that biomechanically, the control of individual joints is limited by the redundant set of muscles that control single or several joints (Lang and Schieber, 2004; Rácz at al., 2012). Studies of neural and neuromuscular architecture of the hand have demonstrated that these do not support fully isolated joint movements (Lemon, 1997; Poliakov and Schieber, 1999; Reilly and Schieber, 2003), and biomechanical constraints appear to result in all muscles being required for full directional control of grip forces (Kutch and Valero-Cuevas, 2011). Additionally, it has been proposed that motor control of the hand joints is organized in a modular way, where several degrees of freedom are organized into functional groups to simplify the control problem (Santello et al., 1998; Tresch et al., 2006).

In the realm of muscle co-actions so called motor synergies were identified to represent structured spatio-temporal patterns of muscle interplay in defined movements (Bernstein, 1967; Santello et al., 2002; Daffershofer et al., 2004; d’Avella et al., 2006; Tresch et al., 2006). Also, studies that have focused on finger joint kinematics of complex hand shapes (Santello et al., 1998; Mason et al., 2001; Daffertshofer et al., 2004), as well as continuous daily-life-day activity (Ingram et al., 2008) found that most variability in the data could be explained by just a few (four to six) characteristic parameters (so called principal components) that indicates a high degree of correlation between the angles of the fingers. These have also been replicated in studies focusing on the key evolutionary ability to produce flint-stone tools (Faisal et al., 2010).

The importance of the hand as our means to interact with the world becomes painfully evident when loss of a hand or hand function occurs. Here neuroprosthetics and robotic hands have rapidly evolved to imitate an unprecedented level of hand-control (Wolpaw and McFarland, 1994; Taylor et al., 2002; Wolpaw and McFarland, 2004; Hochberg et al., 2006; Bitzer and van der Smagt, 2006; Carrozza et al., 2006; Zhou et al., 2007; Kuiken et al., 2007; Steffen et al., 2007; Rothling et al., 2007; Cipriani et al., 2008; Velliste et al, 2008; Lui et al., 2008; Schack and Ritter, 2009; Kuiken et al., 2009; Hochberg et al., 2012; Schröder et al., 2012; Feix et al., 2013; Thomik et al., 2013). Yet, it is still very difficult for people with a lost limb to achieve naturalistic mobility and dexterity by controlling a prosthetic replacement in the same way they would control their own body. This increases the training time to use such neuroprosthetics (up to two years) and results in a low adoption rate after training.

We hypothesize that natural hand movements performed “in-the-wild”, outside artificially construed and highly controlled laboratory tasks contain correlation information that can be used for prediction and reconstruction in the context of prosthetics. We asked subjects to perform everyday tasks such as opening the door, eating, using the phone, etc. The data consists of 15-dimensional time series representing the angles of all the major joints of all the fingers. Advances in experimental methods have increased the availability, amount and quality of high-resolution behavioural data for both humans and animals that can be collected. However, most behavioural studies lack adequate quantitative methods to model behaviour and its variability in a natural manner. Here, we take the view that motor behaviour can be understood by identifying simplicity in the structure of the data, which may reflect upon the underlying control mechanisms. Yet, the analysis of movements and specifically hand movements is complicated by the highly variable nature of behaviour (Faisal et al., 2008). To extract the structure of hand configuration variability data stream we used a probabilistic generative latent variable model (PPCA) of hand configurations for each task.

Part of these results was previously published in the form of abstracts (Belić and Faisal, 2011; Belić and Faisal, 2014).

## 2. Materials and methods

### 2.1. Subjects

Seven adults (two women and five men, average age 24±2 years) with no known history of neurological or musculoskeletal problems, participated in this study following approved ethical guidelines. All subjects were right-handed as determined by the Edinburgh Handedness Inventory (Oldfield, 1971). The experimental procedure used in this experiment was approved by the local ethics committee.

### 2.2. Experiments and data acquisition

We asked subjects to perform 17 different everyday tasks (Figure 1), while capturing their hand movements by using resistive sensors embedded in a previously calibrated CyberGlove I (CyberGlove System LLC, CA, USA). The data glove is made of thin cloth, and its sensors are correlated with corresponding joints of the human hand (Figure 2A). The CyberGlove we used in this study is associated with 18 DOF of the hand. We used data from 15 sensors that consisted of metacarpalphalangeal (MCP) and proximal interphalangeal (PIP) sensors for the four fingers, three stretch sensors between the four fingers, three sensors for the thumb (the carpometacarpal (CMC), MCP and interphalangeal (IP) sensors), and the stretch sensor between the thumb and the palm of the hand. Sensors were sampled continuously at 80 Hz at a resolution of 8 bits per sensor. Subjects completed 10 repetitions for each of the activities, and they always started trials from the same initial position (the hand was placed on the interface device attached to the subject’s belt with the fingers composed together and thumb oriented parallel to the palm). The beginning of each trial was indicated with a sound. The trials were self-paced and the purpose of activities was explained to subjects orally, but they were not instructed about any desired movements for the upcoming trials. After performing the task, the subject then returned his/her hand to the initial position. All programs for data acquisition, visualization and calibration were purpose-developed in C++.

**Figure 1.**
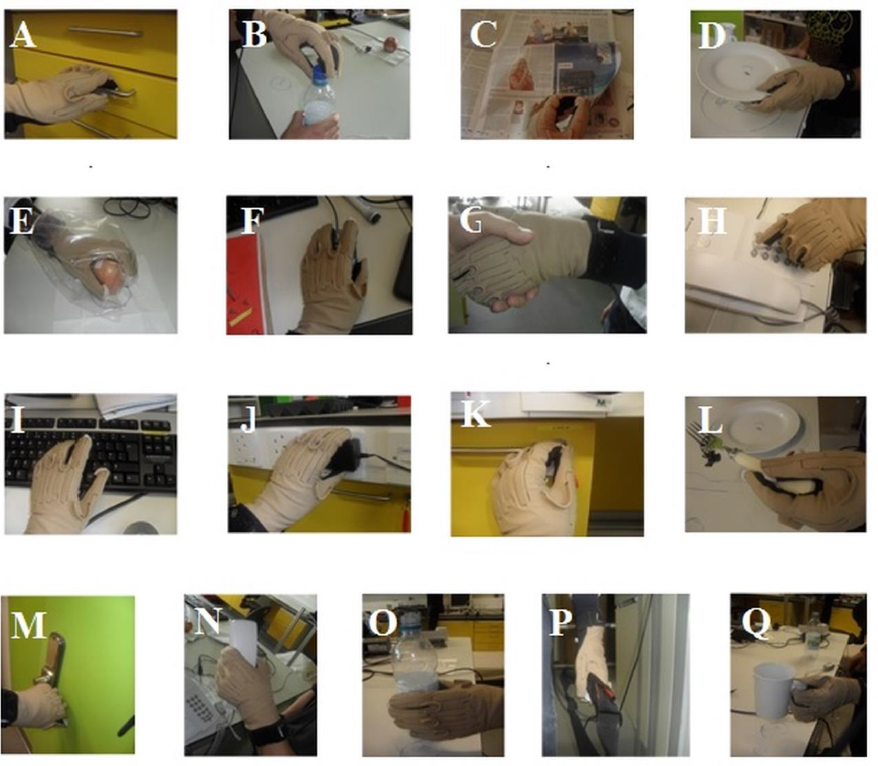
Subjects were involved in 17 different everyday activities. (A) Opening and closing a drawer. (B) Removing a bottle cap (unwinding and winding). (C) Turning the pages of a newspaper (one page in each trial). (D) Picking up a plate, putting it on the marked location, and returning it to the starting position. (E) Eating an apple (subject takes one bite of the apple) and returning the apple to the starting position. (F) Manipulating a mouse in a pre-defined way. (G) Handshaking for a duration of five seconds. (H) Dialling pre-defined numbers on telephone. (I) Typing pre-defined text on a keyboard. (J) Manipulating a plug and returning it to the starting position. (K) Opening a door using a key and returning the key to the starting position. (L) Picking up and putting down an object using a fork. (M) Opening and closing a door using the knob. (N) Picking up a telephone handle. (O) Picking up a plastic bottle, simulating drinking, and returning the bottle to the starting position. (P) Picking up and putting down a bag. (Q) Picking up a glass with a handle, simulating drinking, and returning the glass to the starting position.

### 2.3. Calibration

The output of each CyberGlove sensor is voltage value (raw value) which is dependent on the bending applied to that specific sensor. In order to obtain the outputs in degrees (Figure 2B), it is necessary to determine conversion factor gain and a constant term offset for each of the sensors. This process is called calibration of the CyberGlove. Once the gain and offset are set, output in degrees of the corresponding sensor is given by the following equation:

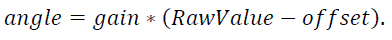

**Figure 2.**
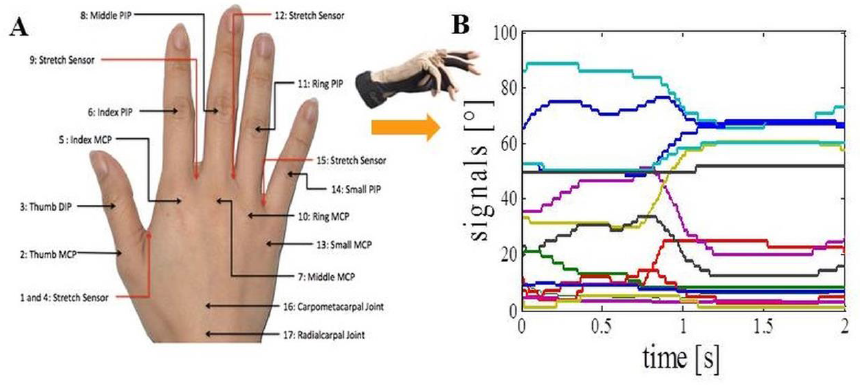
Data acquisition. (A) CyberGlove sensor locations. (B) Calibrated output signals from the CyberGlove for one of the activities.

To calculate the gain and offset we need two different pre-defined angles for each of the sensors and raw values that correspond to them (*RawValue*_1_ and *RawValue*_2_). Gain and offset are calculated by the following formulas:

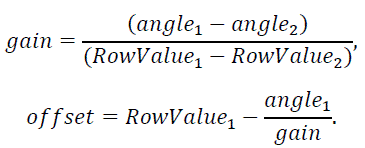

The glove was calibrated for each subject using a five-step procedure that allowed us to determine two different angles (*angle*_1_ and *angle*_2_) for each of the sensors (Figure 3): The first position corresponded to 0° for all glove sensors (Figure 3A).

The second position defined an angle of 90° for all MCP sensors except for the thumb (Figure 3B). The third position determined the abduction angles of 30° between the middle and index finger and between the little and ring finger, an angle of 20° between the ring and middle finger, and an angle of 90° between the index and thumb finger.

**Figure 3.**
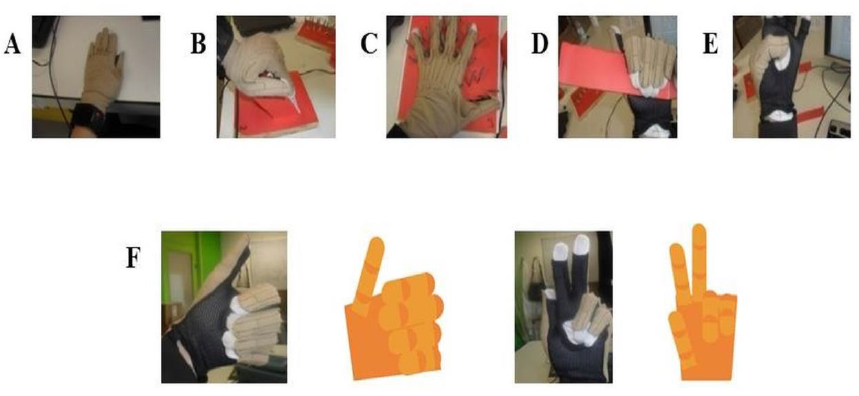
CyberGlove calibration procedures. (A) The position defines *angle_1_* for all CyberGlove sensors. (B) The position defines *angle_2_* for sensors that correspond to MCP joints of the four fingers. (C) The position defines *angle_2_* for abduction sensors. (D) The position defines *angle_2_* for sensors that correspond to PIP joints of the four fingers. (E) The position defines *angle_2_* for sensors that were used to measure the position of the thumb. (F) Examples for real time capturing of finger movements by using a 3D hand model that was developed to further improve accuracy of the calibration procedure.

The fourth position defined an angle of 90° for all PIP sensors except for the thumb.

The fifth position corresponded to the angles for the thumb sensors: CMC (90°), MCP (45°) and IP (90°) sensor.

The calibration procedure was further improved using an online visualization system. In our study, a virtual human hand was rendered in OpenGL. The virtual hand was animated in real-time by data from the glove (Figure 3F). Visualization of data was of great help during both calibration and data acquisition processes. In the case of visually observed deviation between the 3D model and the actual position of the hand, gain and offset were re-determined only for the sensors where deviation was observed. Calibration parameters for each of the subjects had been stored in a separate file and loaded before the experiments started. We also asked subjects, after completing the calibration procedure, to again place their hand in the first position, so we could additionally check eventual discrepancies. The average error across the sensors was 5±2 degrees.

### 2.4. Computational Latent Variable modelling of real-life movements

Collected data from the 15 sensors for each subject and each trial were stored to disk for offline analysis using MatLab (MathWorks, Natick (MA)). Before further analysis, the data is smoothed using a second-order Savitzky-Golay filter with a running window of five data points to remove discontinuities induced by the A/D converter.

Our data space potentially extends over a 15-dimensional space. We performed Principle Component Analysis (PCA) on joint angles in order to estimate real dimensionality of the finger movements during daily activities. PCA reduces the set of correlated variables to a set of non-correlated variables (principle components) (Semmlow, 2001; Bishop 2006). The first principal component contains as much of the variability (as quantified by the variance) in the data as possible, as does each succeeding component for the remaining variability. Therefore, here we used the PCA method to determine the complexity of the finger movements, by measuring how many principal components can explain most of the variability in the data (Faisal et al., 2010). For example, dimensionality reduction techniques can be illustrated by considering the index finger, which has three joints controlled by five muscles. Describing the flexing behaviour of this finger requires *a priori* three values (“dimensions”). For example, in specific movements like making a fist, as we flex one joint of the index finger, we flex the other two joints at the same time in a highly coordinated manner. Thus, we would require in principle a single dimension to describe the configuration of the finger. PCA ignores the temporal structure of movements (in fact the results of PCA will be the same if the data in each trial is randomly shuffled in time). Thus, correct classification relies on the sub-space of finger movement variability alone.

Tipping and Bishop found a probabilistic formulation of PCA by viewing it as a latent variable problem, in which a d-dimensional observed data vector ***x*** can be described in terms of an m dimensional latent vector, ***y***:

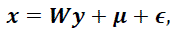

where ***W*** is *d x m* matrix, ***μ*** is the data mean and ***Є*** is an independent Gaussian noise with a diagonal covariance matrix ***I***. The likelihood of observed data vector ***x*** is given as:

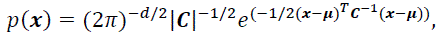

and ***Cov*** is the model covariance matrix given by the following formula:

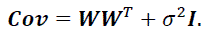

***W*** and *σ* are obtained by iterative maximization of log-likelihood of *p*:

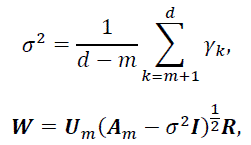

where *γ_k_* are eigenvalues, ***U_m_*** is *d x m* matrix of eigenvectors, ***A_m_*** is diagonal matrix (*m x m*) of eigenvalues, and ***R*** is an arbitrary *m x m* orthogonal rotation matrix (for simplicity ***R*** is usually equal to ***I***).

### 2.5. Measure of manipulative complexity

As a way to quantify manipulative complexity for a given number of PCs, we proposed a universally applicable measure that allowed us to calculate and compare limb movements across different tasks. We refer to it as manipulative complexity *C*, and define the measure by the following formula:

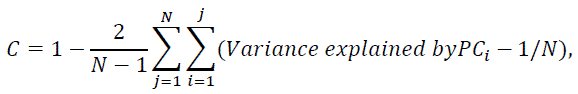

where *N* is the total number of PCs we consider. Our data space extends over a 15-dimensional space, so if all PCs contribute equally that implies *C=1*, and *C=0* if one PC explains all data variability. Our complexity measure compares well with intuitive complexity estimates and it can be thought of as a new assessment measure that is calculated after an objective mathematical analysis. For example, a simple behaviour, e.g. curling and uncurling a hand into a fist, would reveal a single dominant principal component as all 5 fingers (and each finger’s joint) move in a highly correlated manner and therefore *C* would be close to 0. In contrast, a complex behaviour, such as expert typing on a keyboard would reflect more uniform distribution of variances explained by principal components, as each finger moves independently from the others, and so *C* would have a high value.

### 2.6. Task recognition from movement data (Bayesian classification)

Next, we simply predicted a task based on the one with the highest PPCA likelihood by employing Bayesian classifier. In Bayesian statistics there are two important quantities: unobserved parameters *Ω_j_* (*j=1*,…,*17* different activities in our study) and observed data ***x*** (movement data). They are related in the following way:

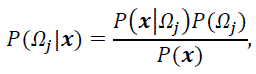

where *P(Ω_j_|****x***), which is termed posterior, represents probability that testing data ***x*** belong to activity *Ω_j_*. Prior, *P(Ω_j_)*, is simply given by the relative frequency of occurrence of each class in the training set and we can ignore it here. Therefore probability of each class, given testing data, is equal to likelihood *P(****x****|Ω_j_)* (probability of seeing the data given the task) that is thoroughly explained in section 2.4.

For training and testing the classifier we used leave-one-repetition (across all actions and all subjects)-out cross-validation.

### 2.7. Missing limb movement reconstruction (Latent variable decoding)

For data reconstruction, firstly we used linear regression to fit the data of missing joints as a function of other joints and expressed results as the average difference between actual and predicted values. Then, we employed the Expectation-Maximization (EM) algorithm for PPCA in order to estimate missing values and at the same time to determine the right subspace dimension. In the EM approach for PPCA, we considered the latent variables ***y_n_*** to be ‘missing’ data and the ‘complete’ data to encompass the observations together with these latent variables (Tipping and Bishop, 1999). The corresponding complete-data log-likelihood is given as:

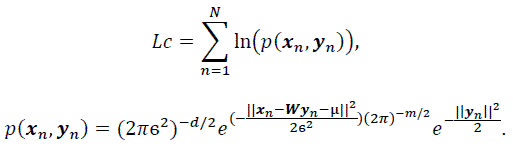

Then we calculated the expectation (E-step) of *L_C_*:

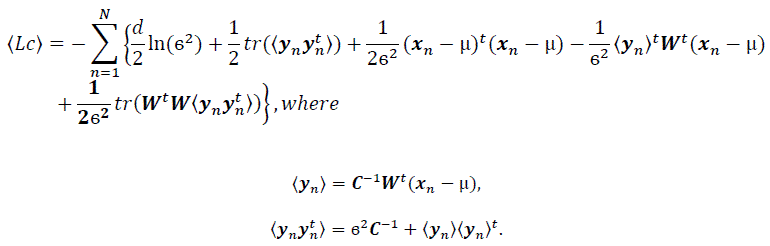

In the M-step, *L_C_* was maximized with respect to ***W*** and *σ^2^*: 

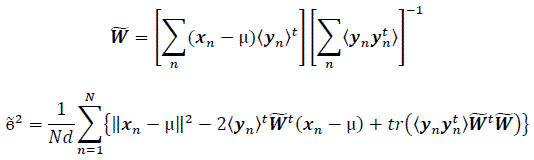

These equations were iterated until the algorithm was judged to have converged.

## 3. Results

### 3.1. Natural hand and finger joint kinematics have a low-dimensional embedding

The structure of natural hand and finger movements in daily-life is characterized by a highly variable nature. Even in the case of handshaking (Figure 1G), which represents one of the most stereotypic actions, basic statistical analysis has revealed vast diversity in angular data for MCP and PIP joints across different subjects (Figure 4A). In this work we first exploited the correlations between MCP and PIP joints for each of the four fingers and we found that correlation coefficients were stronger for little and ring fingers and weaker for middle and index fingers (Figure 4B). Further, correlations between each of the four fingers were highest for the neighbouring fingers and gradually decreased for more distant fingers (Figure 4C). We also used Principal Component Analysis in order to estimate dimensionality of the finger movements across different complex manipulation tasks. Therefore, we used PCA as a measure for the complexity of hand configuration, by measuring the amount of variance in the data displayed by each of the principal components. For example, a simple behaviour such as curling and uncurling the hand would reveal a single dominant PC component, as all finger joints move in a highly correlated manner. In Figure 5A we show the percentage of explained variance versus the number of used principal components for each of the 17 activities. PCA revealed for all tasks that hand motor control restricted hand configurations on a low dimensional subspace of four to five dimensions (which explained 83–96% of the variance in the data), in line with previous data on evolutionary relevant hand behaviour (crafting of flint stone tools, Faisal et al., 2010) and non-annotated long-term statistics of joint velocities (Ingram et al., 2008). These results imply a substantial reduction from the 15 degrees of freedom that were recorded. Some of the activities required more principal components than others to reconstruct the data. For example in Figure 5A we can see that opening a lid on a bottle or manipulating a fork are far more complex activities than hand dialling numbers on a phone. Single PC component explained around 30% less variance in the first case (opening a lid) than in the second (dialling numbers), while that discrepancy was around 10% in the case when we used only the first four PCs to explain variance.

**Figure 4.**
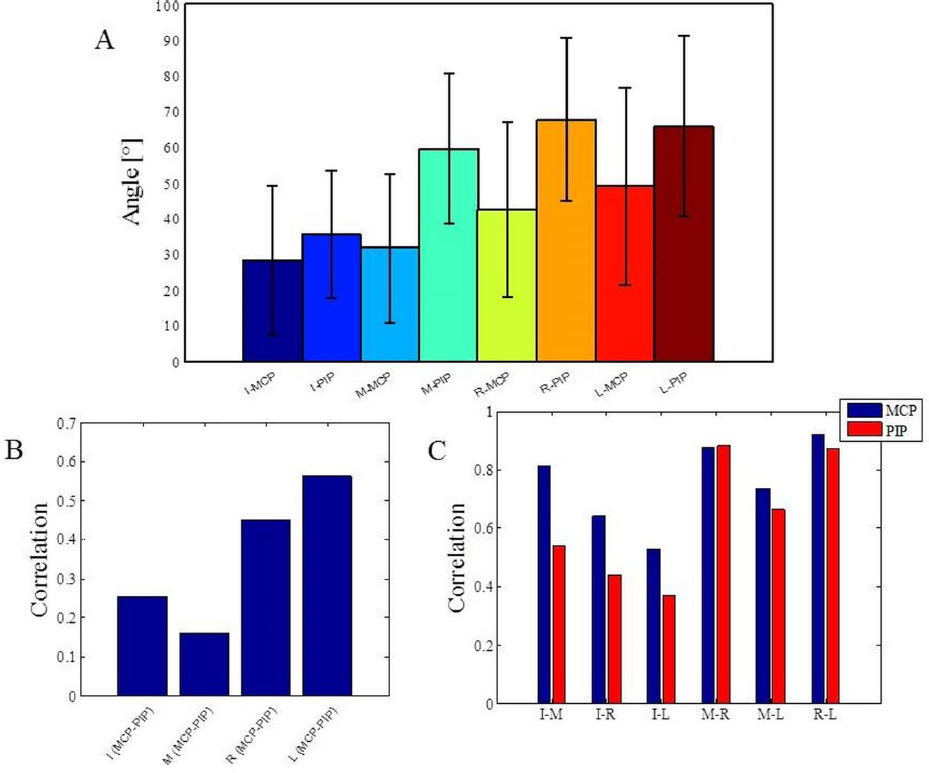
Basic statistics for one of the most stereotypic actions across all subjects and analysis of correlations across all data. (A) Mean and standard error of angular data for MCP and PIP joints in the case of handshaking activity for a duration of five seconds. (B) Correlations between MCP and PIP joints for each of the four fingers. (C) Correlations between each of the four fingers. *I* index finger, *M* middle finger, *R* ring finger, *L* little finger, *MCP* metacarpalphalangeal joint, *PIP* proximal interphalangeal joint.

### 3.2. Measuring the manipulative complexity of activities in daily life

We can visually observe some differences and similarities in the manipulative complexity between the most simple hand movements, during which the individual joints move in a highly correlated manner, and the most complex, where each finger moves independently from the others. Here we proposed a universally applicable measure of manipulative complexity (C) that allows us to measure this quantity across vastly different tasks. Our complexity measure implies that C=1 if all DOF contribute equally (the most complex activities), and C=0 if one DOF explains all DOF (the most simple activities). Results produced are in line with intuitive expectations (opening a lid on a bottle is more complex than operating a door handle) (Figure 5B). Some of the activities in our study also included “grasp like motions” (e.g. operating a door handle, grabbing a bottle or grabbing a bag) that visually would look very similar. Our established complexity measure appeared sensitive enough and was able to differentiate between even those similar looking grasps.

**Figure 5.**
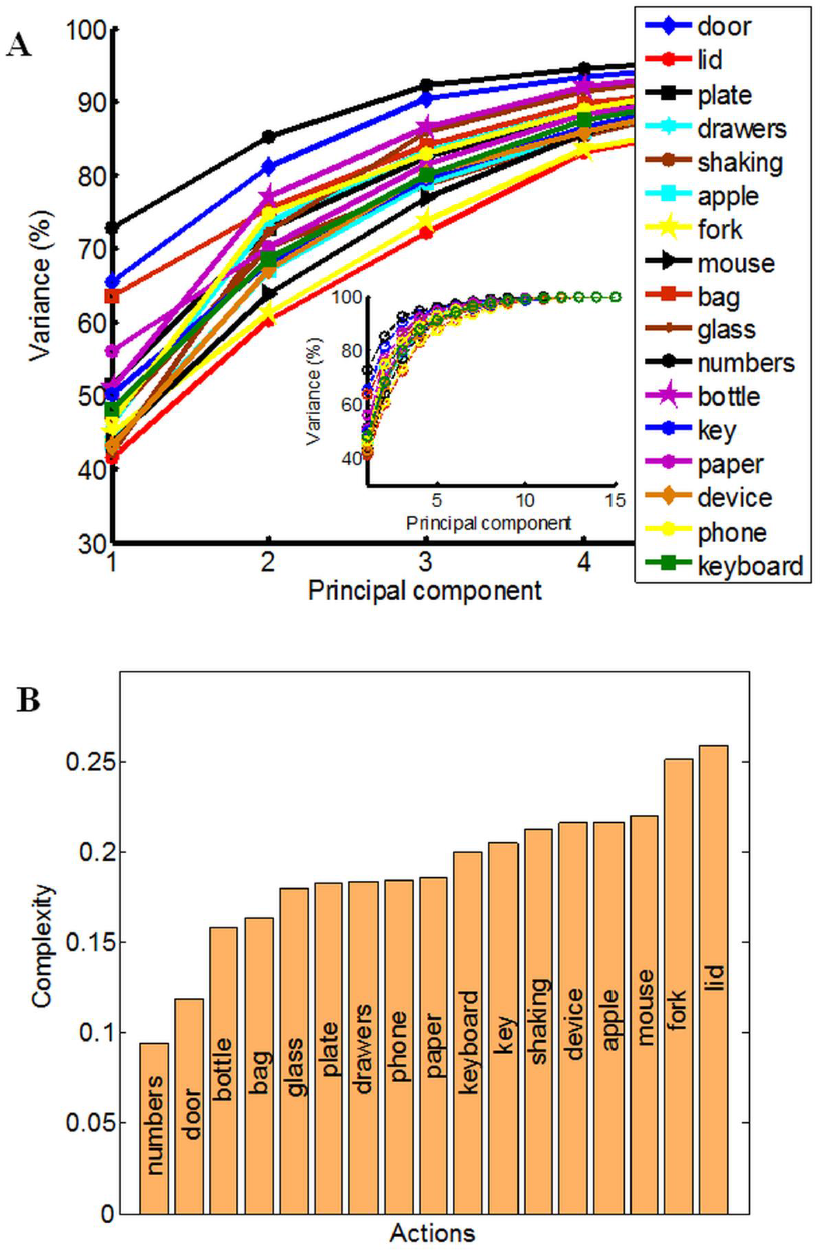
Principal component analysis (PCA) and quantitative measure of manipulative complexity. (A) Curves show the cumulative sum of variance (expressed in percentage) explained by increasing the numbers of principal components separately for each of the 17 daily-life activities. The x-axis corresponds to the number of PCs; the y-axis shows the percentage of the variance of the finger movements explained by the respective number of PCs. (B) Our proposed quantitative measure of manipulative complexity (manipulative complexity is maximal (equal to 1) if all DOF contribute equally, and minimal (equal to 0) if one DOF explains all DOF).

### 3.3. Prediction of hand movements from initial movement data

Further, we wanted to see how different subspaces influence success of classification for different tasks. To deal with this, we used Bayesian PPCA. PPCA has been considered as a mechanism for probabilistic dimension reduction or as a variable-complexity predictive density model (Tipping and Bishop, 1999) and correct classification relies on the subspace of finger movement variability alone. Figure 6A illustrates the success of classification with reference to the number of PPCA components. Therefore, by using only the first few PPCA components in the classification process we can get very high classification success. For example, using the first four PPCA components the success of classification was 89.91% (across all tasks, classification performance was 96.63% using all 15 PPCA components). Importantly, in Figure 5 one could see that extracted subspaces appeared to be task-dependant, which suggests that besides simplification, synergies might have a role in a task-optimal control as well. If specific tasks can engage specific motor control strategies, then we should be able to make a conclusion regarding the task by observing some early portions of finger data. Indeed, the classification performance, presented in Figure 6B, was a few times higher than the chance performance (marked with red line) for only an initial portion of the finger configuration samples of each task. Within the first 1000ms from the initial hand position, which was identical for every action, it appeared that hand shape already configures itself to a specific task and we were able to quickly predict intended action. Vertical lines represent average duration for each of 17 activities.

**Figure 6.**
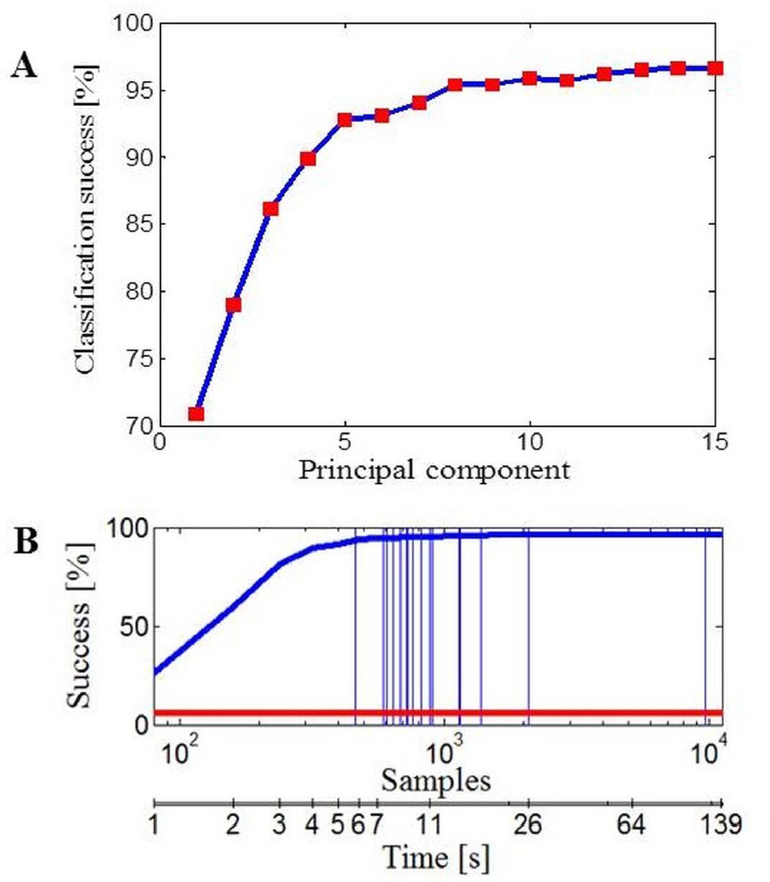
Data classification. (A) Classification performance with reference to the number of PPCA components. (B) Classification performance with reference to the number of data samples taken (duration of activity). Performance by chance is marked with the red line and vertical lines represent the average number of data samples (duration) for each of 17 activities.

### 3.4. Reconstruction of missing limbs’ movements by decoding movements of remaining limbs

Next, we investigated the predictability of a subset of joint movements in respect of the movements of other joints. Or in other words, if part of a limb is missing, how well can we predict what those missing parts should be doing by only observing the intact, remaining limb parts. This is of fundamental interest in prosthetic control. We focused particularly on cases where data had been acquired with sensors that measure the bending around the MCP or the PIP joints of the four missing fingers. First, we applied linear regression in the case of missing values from the MCP joints (Figure 7A) and the PIP joints (Figure 7B) for each of the four fingers separately. The error we got, measured as absolute difference between predicted and actual joint values and averaged across all tasks, showed the best linear predictability for the middle and ring fingers in both examined cases. Overall predictability rate was high regarding movement range (90 °) for each of the considered joints and variability of tasks. Then, we applied an EM algorithm for PPCA to infer the un-observed, invisible joints in the case of missing data from the MCP sensors. Figure 7C shows obtained results with reference to the number of PPCA components. Here the best results were also acquired for the middle and ring fingers. The error was the highest in the case when just one PPCA component was used and then started to decrease (up to a number around 8 PPCA). Generally, these results could help us to improve the method of designing prosthetic controllers that are driven by intact limb parts and support neuroprosthetic controllers in refining the decoding of action intention of users.

**Figure 7.**
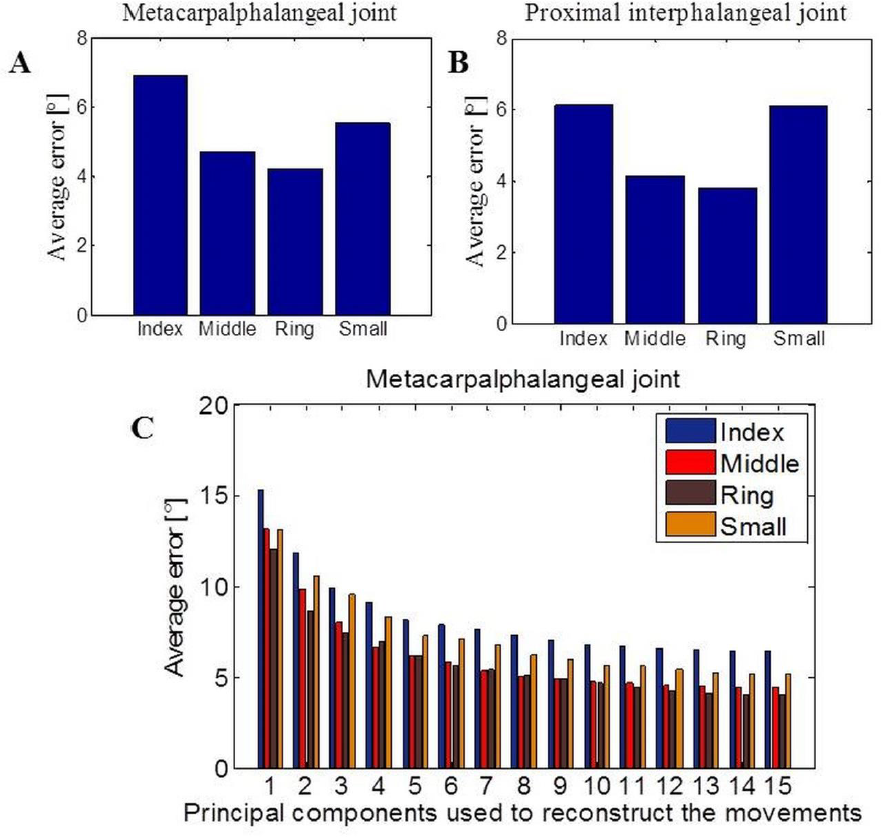
Data reconstruction. (A) Average error after linear reconstruction in the case when data from MCP sensors of the four fingers were missing. (B) Average error after linear reconstruction in the case when data from PIP sensors of the four fingers were missing. (C) Results of data reconstruction by using PPCA with reference to the different number of PC components used.

## 4. Discussion

We analysed natural movements from the seven subjects who were behaving spontaneously while performing 17 different everyday activities. We have four key findings that we will discuss individually in more detail as follows: 1. Regarding activities of daily living, we confirmed that hand control is low-dimensional, i.e. four to five PCs explained 80–90% of the variability in the movement data. 2. We established a universally applicable measure of manipulative complexity that allowed us to measure this quantity across vastly different tasks. Our findings are in line with intuitive expectations (opening a lid on a bottle is more complex than hand dialling numbers) and are sensitive enough to differentiate between similar looking interactions (e.g. operating a door handle is less complex than grabbing a bottle). 3. We discovered that within the first 1000ms of an action the hand shape already configures itself to vastly different tasks, enabling us to reliably predict the action intention. 4. We suggest how the statistics of natural finger movements paired with Bayesian latent variable model can be used to infer the movements of missing limbs from the movements of the existing limbs to control for example, a prosthetic device.

In many everyday activities we move our fingers in a highly correlated manner. Therefore, it has been proposed that control of human hand movements is organized in a way that comprises coupling of several DOF into functional groups. The opinion that motor synergies lie behind manual actions has been supported by several studies (Santello et al., 1998; Santello et al., 2002; Daffershofer et al., 2004; d’Avella et al., 2006; Tresch et al., 2006; Ingram et al., 2008; Faisal et al., 2010; Jarrasse et al., 2014). The most common interpretation is in terms of simplifying the strategy that the central nervous system might undertake. Studies that have investigated hand configurations during reaching and grasping movements (Santelo et al., 1998; Mason et al., 2001) reported that 90% of the variance in hand configurations could be explained by only three principal components. In our study PCA analysis revealed that in 17 daily activities hand configurations operated on low-dimensional subspace (four to five dimensions) as well, which is also in line with previous data on evolutionary relevant hand behaviour (crafting of flintstone tools) (Faisal et al., 2010) and non-annotated long-term statistics of joint velocities (Ingram et al., 2008). These finding supports the view that the motor cortex organizes behaviour in a low-dimensional manner to avoid the curse of dimensionality in terms of computational complexity. We also found numerical differences in the number of principle components required to explain a given amount of variability in hand configurations across each of the tasks. Similar conclusions were obtained in the case of a small number of much simpler manipulation tasks (Todorov and Ghahramani, 2004; Bläsing et al., 2013).

Our manipulative complexity measure, established for the first time, gave us a chance to quantify the complexity of the movements across a high number of different activities. This was very important in that some of the activities that look highly similar (grasp like motions such as operating a door handle, grabbing a bottle or grabbing a bag) apparently had different values of complexity. Those findings demonstrated also that our complexity measure is sensitive enough to differentiate between similar looking interactions. The highest value of complexity had tasks of opening a lid on a bottle or manipulating a fork, and the lowest had tasks of dialling numbers on a phone or opening a door using the door knob. Results produced are in line with intuitive expectations regarding the fact that in the first two cases one is expected to have high engagement of the thumb that is the most individuated (Häger-Ross and Schieber, 2000). In the case of typing numbers, most of the subjects used their index finger while their other fingers created some form of fist, and in case of opening the door our fingers move in a highly correlated manner. Here we compare structures of complex dynamic hand manipulations, while some other studies (Feix et al., 2009; Feix et al., 2013) have presented a successful methodology for measuring and evaluating the capability of artificial hands to produce 31 different human-like grasp postures.

Further we employed Bayesian PPCA on the behavioural data in order to analyse the structure of variability within it. Variability is ubiquitous in the nervous system and it has been observed in actions even when external conditions are kept constant (Faisal et al., 2008). In this paper we take the view that the hand configuration variability may contain significant information about the task being performed. Our approach yields an effective assessment of the tasks that subjects were involved with. The Bayesian PPCA reveals that the finger movement correlations are so structured that we can obtain very high classification success by taking only first few principal components. Regarding motor control, it has been suggested that structural learning (Braun et al., 2009) may reduce the dimensionality of the control problem that the learning organism has to search in order to adapt to a new task. Our results are in line with this concept and suggest the hypothesis that the brain can engage many sets of motor controllers, which are selected based on specific tasks, and which also orchestrate resulting actions in overall behaviour and produce movement variability in characteristic sub-spaces. Next we thought that, if the hypothesis is true, we should be able to infer the task the hand is engaged in by observing some initial portion of the finger movement data. Crucially, observing only the initial portion of hand configurations (from our identical starting position) was sufficient to characterize the entire hand task, and the classification performance we obtained was a few times higher than chance performance.

A common approach in design of Neuroprosthetics is to construct body parts that can be controlled with the same functionality as natural limbs. Using a reduced set of basic functions to construct internal neural representation could be essential from an optimal control perspective (Poggio and Bizzi, 2004) and applied to Neuroprosthetics control (Thomik, et al., 2013). Our linear predictability of the missing joints based on movements of other finger joints gave good results. The best results were achieved for middle and ring fingers showing that they are the least individuated. This is in line with the previous research (Häger-Ross and Schieber, 2000). Further, the PPCA algorithm for missing data revealed that using more than eight PPCA components does not lead to any significant improvement. In this study we perform action recognition and reconstruction of missing finger trajectories using the current positions of other functional finger joints by simply requiring – in principle – the user to act out with his functional fingers an intended task. Such finger motion can be realized with cheap wearable wireless sensors (Gavriel and Faisal, 2013) and we can reconstruct the natural behaviour of users without the need for expensive, training intensive, non-invasive or invasive electrophysiological interfaces. Consequently, unlike common approaches that require the user to learn to use the technology, the technology interprets the natural behaviour of users (Abbott and Faisal, 2011). Thus, the neuronal and biomechanically imposed correlation structure of hand-finger can be exploited to build smart, sensitive Neuroprosthetics controllers that infer the task “at hand” based on the movements of the remaining joints.

Dexterous object manipulation is conditioned by the continuous interactions between the body and the environment and engages multiple sensory systems. Vison can provide essential information for controlling hand kinematics in the cases when object are fully visible. Human manipulation involves also tactile signals from different types of mechanoreceptors in the hand that allows humans to easily hold a very wide range of objects with different properties without crushing or dropping them (Johansson and Flanagan, 2009). Tactile sensing provides also critical information in avoiding slipping as crucial precondition to successfully manipulate an object, what is most apparent in people with impaired tactile signals. When finger contact with the desired object is made, we start to increase the grasp force to the optimal level, using both our prior knowledge about the object and information from the tactile sensors of the fingers gathered during the interaction (Johansson and Flanagan, 2008; Romano et al., 2011). Corrective actions are applied to different frictional conditions in order to provide an optimal grip force that is normally 10–40% greater than the minimum required to prevent slips (Johansson and Flanagan, 2008). Consequently, future Neuroprosthetics should provide reliable user’s intention decoding as well as optimal sensory feedback (Berg et al., 2013; Raspopovic et al., 2014). Therefore, looking into hand kinematics as an important aspect of the hand capabilities represents just one approach that forms the basis for future studies. Further inclusion of other parameters that are of relevance and investigating their influence on manipulative complexity will provide a more complete analysis.

## 5. Acknowledgments

JJB was supported by the IASTE scholarship of the British Council. AAF acknowledges the support of the Human Frontiers Program Grant (HFSP RPG00022/2012).

